# An integrated landscape of protein expression in human cancer

**DOI:** 10.1101/665968

**Authors:** Andrew F. Jarnuczak, Hanna Najgebauer, Mitra Barzine, Deepti J. Kundu, Fatemeh Ghavidel, Yasset Perez-Riverol, Irene Papatheodorou, Alvis Brazma, Juan Antonio Vizcaíno

**Author notes:** Corresponding authors: Dr. Andrew F. Jarnuczak; European Molecular Biology Laboratory, European Bioinformatics Institute (EMBL-EBI), Wellcome Trust Genome Campus, Hinxton, Cambridge, CB10 1SD, UK.; Dr. Juan Antonio Vizcaíno; European Molecular Biology Laboratory, European Bioinformatics Institute (EMBL-EBI), Wellcome Trust Genome Campus, Hinxton, Cambridge, CB10 1SD, UK.

## Abstract

Using public proteomics datasets, mostly available through the PRIDE database, we assembled a proteomics resource for 191 cancer cell lines and 246 clinical tumour samples, across 13 cancer lineages. We found that baseline protein abundance in cell lines was generally representative of tumours. However, when considering differences in protein expression between tumour subtypes, as exemplified in the breast lineage, many of these changes were no longer recapitulated in the cell line models. Integration of proteomics and transcriptomics data suggested that the level of transcriptional control in cell lines changed significantly depending on their lineage. Additionally, in agreement with previous studies, variation in mRNA levels was often a poor predictor of changes in protein abundance. To our knowledge, this work constitutes the first meta-analysis study including cancer-related proteomics datasets. We anticipate this aggregated dataset will be of significant aid to future studies requiring a reference to baseline protein expression in cancer.

## INTRODUCTION

Cancer cell lines are powerful models often used in place of primary cells to study for instance the molecular mechanisms of the disease. Cell lines derived from patients provide an inexpensive source of pure cell populations and are easy to manipulate and characterize. Although they usually retain driver mutations, cell lines often contain ‘additional’ genomic aberrations not present in tumours. They also lack the tumour microenvironment interactions and can undergo divergent evolution during long-term cell culture^1–3^. Hence, in order to capture the patient’s tumour biology, it is therefore often more desirable to study primary cells or tissue biopsies.

Thanks to a number of International initiatives and large-scale studies, a variety of omics approaches have been employed to characterise both tumour samples and cell lines at the molecular level. Among them, The Cancer Genome Atlas (TCGA) used next-generation sequencing (NGS) and reverse-phase protein arrays (RPPAs) to generate genome and expression landscapes in approximately 10,000 tumour specimens^4^. The Cancer Cell Line Encyclopedia (CCLE) consortium measured DNA copy-number, mutations and gene expression in 1,072 human cancer cell lines^5^. Also in this context, the Genomics of Drug Sensitivity in Cancer (GDSC) project provided genomic and gene expression data for over 1,000 cell lines combined with cell line sensitivity measurements related to a wide range of anti-cancer therapeutics^6^.

While the majority of the available data is based on NGS technologies measuring DNA and RNA-level alterations in cancer, proteins are however most often the functional molecules, providing a link between genotype and the phenotype. Proteins are also the targets of many drugs and can be a source of novel biomarkers. Mass spectrometry (MS) is the main proteomics technology, capable of providing system-wide measurements of protein expression, post-translational modifications and/or protein-protein interactions, among other pieces of information^7, 8^. Accordingly, cancer cell lines proteomes have been characterised in a number of MS-based studies^9–17^. However, due to technological limitations, these and other currently publicly available proteomics datasets are usually smaller in scope in comparison to analogous genomics and transcriptomics studies. Therefore, although they can routinely cover thousands of gene products, the datasets are usually limited to tens of samples. MS-based protein expression information in tumour specimens has also been obtained, for example through the work of the Clinical Proteomic Tumour Analysis Consortium (CPTAC)^18, 19^ or by other independent groups^20–22^. Additionally, RPPA approaches can be used to characterise a usually much smaller number of proteins^5, 23^.

Importantly, the MS data underpinning these efforts is now routinely made freely available in the public domain, an opposite situation to the state-of-the-art just a few years ago. Particularly, the PRIDE database^24^ is the world-leading resource as part of the ProteomeXchange Consortium, storing raw files, processed results and the related metadata coming from thousands of original datasets. Public datasets stored in PRIDE, together with existing ones in other open proteomics repositories (e.g. the CPTAC portal^25^ or MassIVE^26^) present an opportunity to be systematically reanalysed and integrated, in order to confirm the original results, potentially obtain new insights and be able to answer biologically relevant questions orthogonal to those posed in the original studies. Such integrative meta-analyses have already been successfully employed in different omics data types, especially in genomics and transcriptomics^27^. For example, Lukk *et al.*^28^ integrated thousands of microarray files to compile a map of human gene expression. In metabolomics, Reznik *et al.*^29^ used MS data from eleven studies to measure the extent of metabolic variation across tumours. A similar trend is starting to be observed in proteomics, where reuse of public datasets is becoming increasingly popular, with multiple applications^30, 31^. Some examples where joint reanalysis of large public datasets has been performed involved the creation of comprehensive maps of the human proteome^32^ and of human protein complexes^33^, or the characterisation of the functional human phosphoproteome^34^.

However, to our knowledge no previous studies have attempted to reanalyse and integrate quantitative proteomics datasets in order to provide a global reference map of protein expression in cancer. Here, we provide a reference resource of protein expression across different types of primary tumours and the corresponding cell line models (246 clinical tumour samples and 191 cancer cell lines), using public proteomics datasets as the base.

## RESULTS

### A catalogue of cancer cell lines and primary tumour proteomes

We selected, manually curated and re-annotated 7,171 MS runs coming from 11 large-scale quantitative cancer related proteomics studies (Table 1) (**Figure** 1A). The combined analysis yielded an aggregated dataset of protein expression in 191 cancer cell lines and 246 clinical tumour samples (**Figure** 1B). In addition, 35 non-malignant tissues, present in the original publications, were also included in the combined dataset. Cell line samples originating from 13 different tissue-origins were included: blood, bone, brain, breast, cervix, colorectal/large intestine, kidney, liver, lung, lymph node, ovarian, prostate and skin. The patient-derived samples came from breast, colorectal, ovarian and prostate tumours, and from breast to lymph node metastases. Lineage annotation of the cell lines and tumour samples is included in Supplementary Table 1. Overall, over 173 M spectra were reanalysed using MaxQuant (MQ)^35^. The resulting aggregated dataset covered 15,443 gene products with at least one unique (unambiguous) peptide evidence, which corresponded to a 67.8% coverage of the entire UniProt reference human proteome (**Figure** 1**C**). Since quantitative proteomics data originating from different studies can be heterogeneous and likely to contain batch effects, we developed and benchmarked a procedure to integrate appropriately the quantification data (Methods section, see Supplementary Methods for detailed description). Based on that, we obtained quantification values for an average of 6,593 proteins per cancer cell line and 5,371 proteins per tumour type.

**Figure 1.**
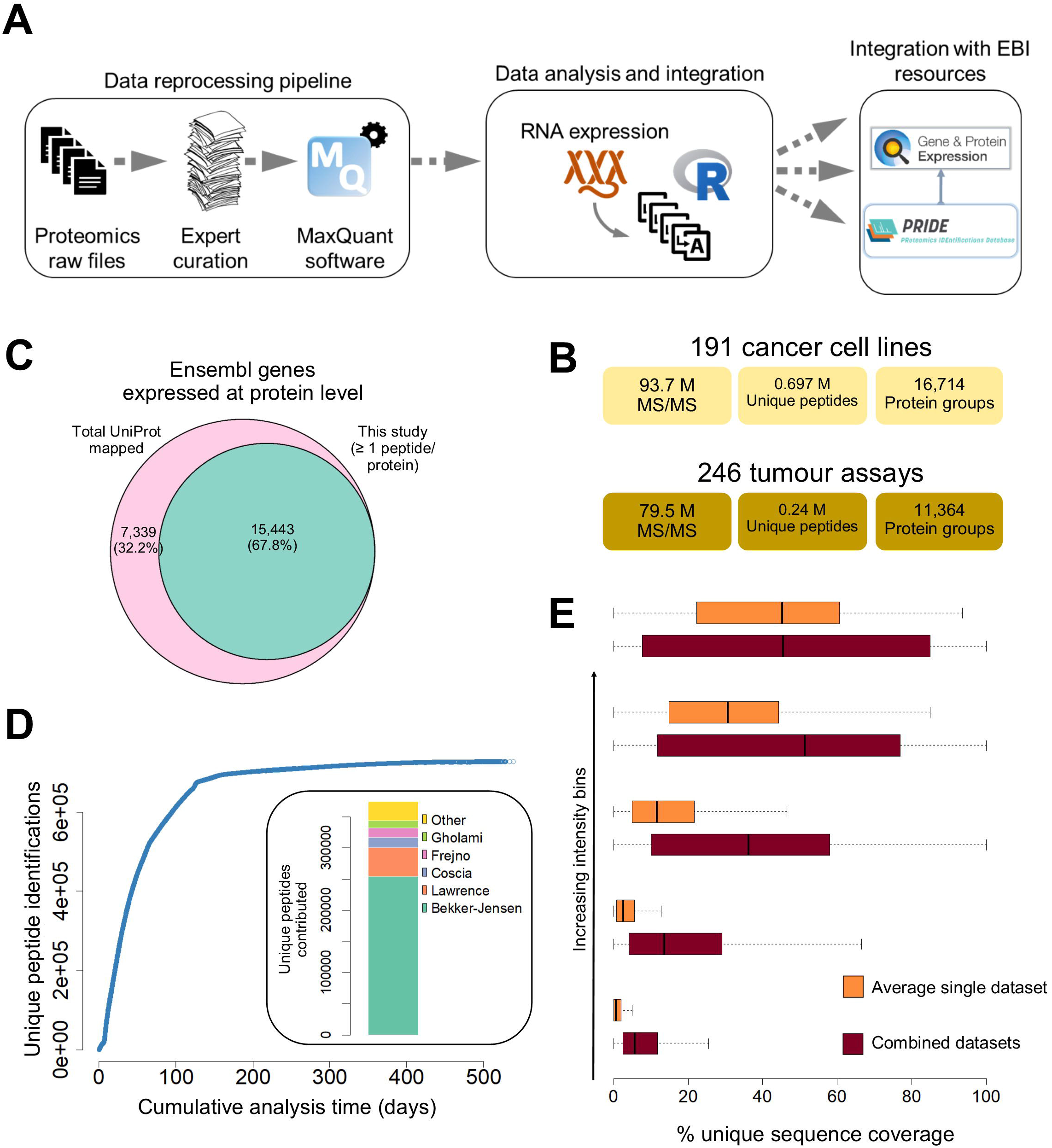
A) Overview of the study design and the data reanalysis pipeline. B) Summary of the total number of MS/MS spectra, number of unique peptides and protein groups identified in the cell line and tumour datasets. C) Overlap between Ensembl protein-coding genes identified and all theoretical genes annotated in UniProt. Only protein groups identified by at least one unique peptide were included. D) Plot showing the cumulative MS analysis time versus the cumulative number of unique peptide identifications obtained. Inset: barplot showing the proportion of peptides uniquely identified in each individual dataset. E) Distribution of protein sequence coverage in the cell line data, stratified into bins of increasing protein intensity, in the combined dataset (brown boxplots) and an average calculated across 7 individual datasets (orange boxplots).

**Table 1.**
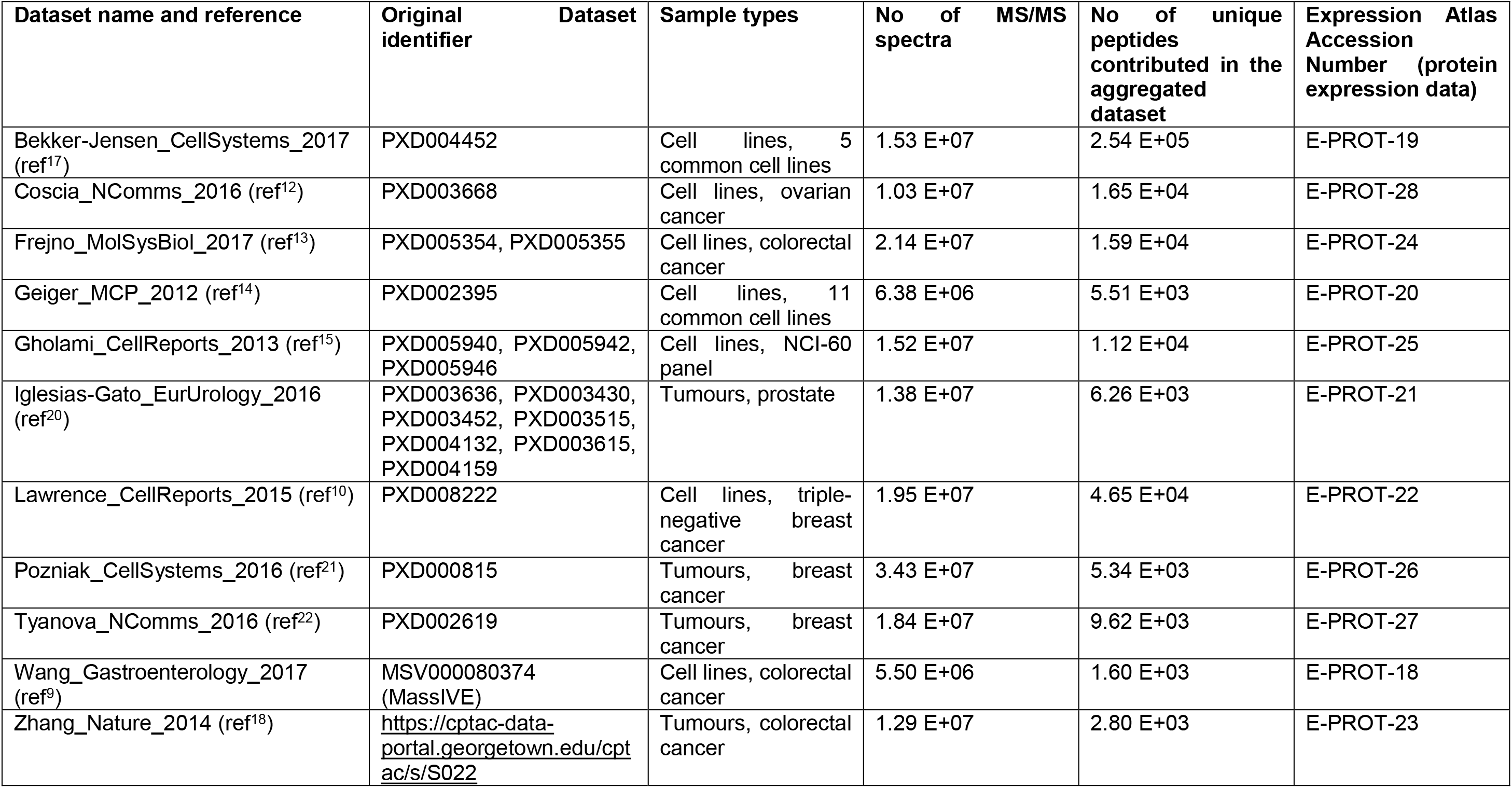
List of the public datasets used in this study. Most of the datasets were obtained from the PRIDE database^24^. The remaining two datasets were obtained from MassIVE^26^ and the CPTAC data portal. The mapping between sample names and their lineage/cancer type and study of origin are listed in Supplementary Table 1.

| Dataset name and reference | Original Dataset identifier | Sample types | No of MS/MS spectra | No of unique peptides contributed in the aggregated dataset | Expression Atlas Accession Number (protein expression data) |
| --- | --- | --- | --- | --- | --- |
| Bekker-Jensen_CellSystems_2017 (ref<sup>17</sup>) | PXD004452 | Cell lines, 5 common cell lines | 1.53 E+07 | 2.54 E+05 | E-PROT-19 |
| Coscia_NComms_2016 (ref<sup>12</sup>) | PXD003668 | Cell lines, ovarian cancer | 1.03 E+07 | 1.65 E+04 | E-PROT-28 |
| Frejno_MolSysBiol_2017 (ref<sup>13</sup>) | PXD005354, PXD005355 | Cell lines, colorectal cancer | 2.14 E+07 | 1.59 E+04 | E-PROT-24 |
| Geiger_MCP_2012 (ref<sup>14</sup>) | PXD002395 | Cell lines, 11 common cell lines | 6.38 E+06 | 5.51 E+03 | E-PROT-20 |
| Gholami_CellReports_2013 (ref<sup>15</sup>) | PXD005940, PXD005942, PXD005946 | Cell lines, NCI-60 panel | 1.52 E+07 | 1.12 E+04 | E-PROT-25 |
| Iglesias-Gato_EurUrology_2016 (ref<sup>20</sup>) | PXD003636, PXD003430, PXD003452, PXD003515, PXD004132, PXD003615, PXD004159 | Tumours, prostate | 1.38 E+07 | 6.26 E+03 | E-PROT-21 |
| Lawrence_CellReports_2015 (ref<sup>10</sup>) | PXD008222 | Cell lines, triple-negative breast cancer | 1.95 E+07 | 4.65 E+04 | E-PROT-22 |
| Pozniak_CellSystems_2016 (ref<sup>21</sup>) | PXD000815 | Tumours, breast cancer | 3.43 E+07 | 5.34 E+03 | E-PROT-26 |
| Tyanova_NComms_2016 (ref<sup>22</sup>) | PXD002619 | Tumours, breast cancer | 1.84 E+07 | 9.62 E+03 | E-PROT-27 |
| Wang_Gastroenterology_2017 (ref<sup>9</sup>) | MSV000080374 (MassIVE) | Cell lines, colorectal cancer | 5.50 E+06 | 1.60 E+03 | E-PROT-18 |
| Zhang_Nature_2014 (ref<sup>18</sup>) | https://cptac-data-portal.georgetown.edu/cptac/s/S022 | Tumours, colorectal cancer | 1.29 E+07 | 2.80 E+03 | E-PROT-23 |

From the information available in the raw files we inferred that it would take over 538 days of mass spectrometer time to repeat all of the original experiments (**Figure** 1D), ascertaining the potentially huge benefit to performing the *in silico* data reanalysis. In addition, the aggregation of individual datasets increased the global proteome coverage. In this study, each dataset contributed was quite heterogeneous and ranged between 1,600 and 250,000 unique peptide identifications to the aggregated dataset (**Figure** 1D inset), in parallel increasing the confidence and robustness in the protein identification and quantification analyses. This was particularly true for low abundance proteins, where the average protein sequence coverage in the aggregated dataset was typically higher than the sequence coverage found in individual datasets (**Figure** 1E).

The overall results of the study have been made publicly available in two EMBL-EBI resources: Expression Atlas (EA)^36^ and the PRIDE database^24^. Expression Atlas provides the expression values for each dataset in a separate track (E-PROT-18 to E-PROT-28, Table 1, **Supplementary Figure 1**). PRIDE dataset identifier PXD013455 contains all the raw data, MQ intermediate files and the combined proteomics analysis results.

### Tumour-specific peptide signatures are enriched in receptor activity regulators

Our first objective was to use the peptides identified in the aggregated dataset to determine whether any clear differences, independent of the tissue of origin and/or the lineage, existed between the tumour and the cell line proteomes. On average, 6,208 proteins were detected in the majority of tumours of any given type (meaning in ≥ 50% of samples of that group) and 7,401 proteins in cell line data (Supplementary **Figure** 2).

When we compared the peptides identified across all cancer cell lines versus all the tumours, we observed that only a small fraction was identified exclusively in tumour data. Out of 711,352 peptides, 33,045 (4.6%) were present only in tumours, constituting a *tumour-specific peptide set*. In contrast, a much larger proportion of peptides was identified only in cell lines (66.0%). This was expected as some of the cell line studies employed extensive fractionation protocols or used multiple digestion enzymes (for example, in dataset from ref^17^ the authors used four enzymes to obtain a deep proteome of HeLa, Table 1). From the *tumour-specific peptide set*, only peptides that uniquely matched to a protein sequence were retained, therefore enabling the unambiguous identification of the corresponding proteins. These 9,907 peptides mapped to 330 proteins for which no identification evidence was found in any of the cell lines. This is in our view an interesting finding given that the sequence coverage of the cell line datasets was much larger. Next, gene ontology (GO) enrichment analysis of this protein set was performed using GOrilla^37^ and REVIGO^38^ (Methods section), revealing that the tu mour-specific proteins were most significantly enriched for biological processes associated with *regulation of signalling receptor activity* (GO:0010469). Other terms, significantly enriched at a FDR (False Discovery Rate) p-value < 0.05 level, included *keratinization* (GO:0031424), *G protein-coupled receptor signalling pathway* (GO:0007186), *positive regulation of leukocyte chemotaxis* (GO:0002690), *humoral immune response* (GO:0006959) and *response to bacterium* (GO:0009617) (Figure ***2***) (Supplementary Table 2). In addition, this protein set was enriched in *extracellular space* related cellular component GO terms (Supplementary Table 3), suggesting that tumour-specific proteins could be involved in secretion. No tumour-specific proteins were consistently detected across all the tumour samples. However, ovarian, colorectal and prostate tumours showed lineage specific expression, as highlighted in Figure ***2***B. After performing a pathway enrichment analysis using Reactome^39^ (Methods section) we found that the 10 proteins specific to the ovarian tumour samples were immunoglobulins enriched in elements of the *CD22 mediated BCR regulation pathway* (Reactome pathway identifier R-HSA-5690714, FDR p-value= 2.89 E-15).

**Figure 2.**
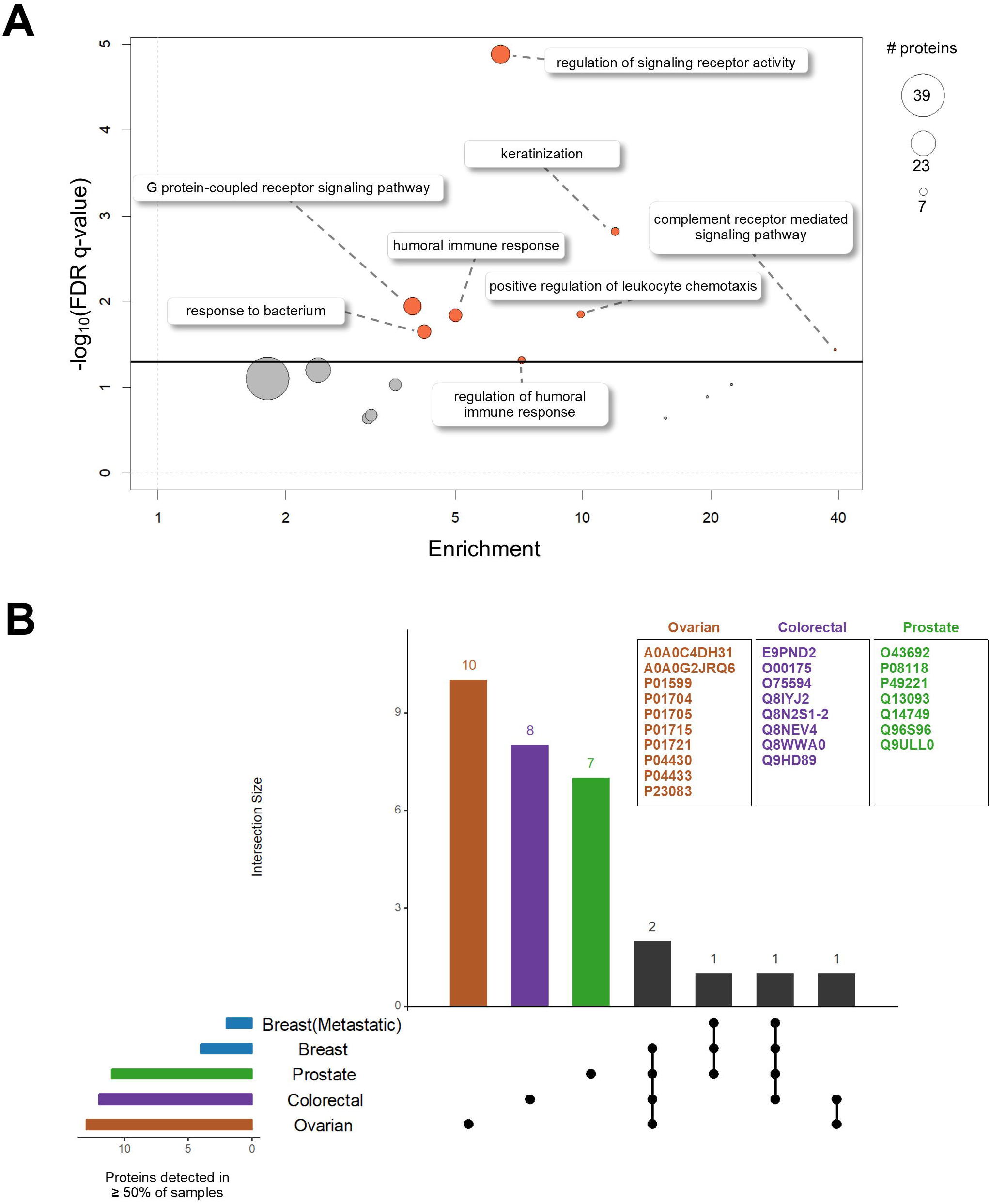
A) Scatterplot summarising the enrichment analysis of biological process GO terms for the 300 proteins detected only in tumour samples. The x-axis represents the enrichment score, the y-axis represents the enrichment p-value corrected for multiple testing using the Benjamini and Hochberg method. The size of the circles corresponds to the number of proteins associated within a given category. B) Counts out of the 330 tumour-specific proteins detected in the majority of samples (i.e. *≥* 50% of samples) across different tumour types. Colour coded horizontal bars show the total number of proteins detected in the majority of samples in each tumour type. Black circles indicate the intersections. Horizontal bars in the histogram show the number of proteins in the corresponding intersection. Proteins specific (as UniProt protein identifiers) to each tumour type are listed in the text boxes.

Taken together, the analysis highlights biological processes specific to tumour samples, which would be difficult to detect and study in cell lines, since expression of proteins associated with those pathways does not seem to be detectable in the cell lines considered here.

### Evaluating cell lines as tumour models based on protein expression

In order to go beyond simple presence/absence qualitative measurements, we generated two matrices containing normalised protein expression measurements across cell lines and tumour samples (Methods and Supplementary Methods). We merged the two by cross-referencing the leading razor protein identifiers, which resulted in a complete matrix containing profiles for 4,476 proteins. We used these quantitative values to investigate the similarity between the samples.

Compared with the tumour tissues, cell lines showed similar levels of variability in protein expression. This was established using a coefficient of variation (CV) calculated between different cell lines and tumours within a given tumour type (i.e. reflecting cancer lineage sample-to-sample variability) (Supplementary **Figure** 3A). The median CV was below 56% in all cases and in general, the majority of proteins had CV values below 70% in both cell lines and tumours. Unsurprisingly, the variability between biological replicates (assessed in cell lines only) was lower than between samples from a different biological origin (median CV = 43%, Supplementary **Figure** 3B).

To investigate the level of concordance in protein expression between endogenous tumour cells and their corresponding cell line models, Pearson correlation coefficient (r*_p_*) values were calculated between each tumour sample and all available cell lines matching that lineage. This was done for three lineages: colorectal (**Figure** 3A), breast (**Figure** 3B), and ovarian samples (**Figure** 3C). Overall, molecular profiles appeared similar between samples, as reflected by relatively high r*_p_*, ranging from 0.58 to 0.83 (p-values < 2.2 E-16) (**Figure** 3A-C) and the median r*_p_* of 0.73, indicating that cell lines can generally represent the baseline protein expression present in tumours. Interestingly, in case of colorectal (**Figure** 3A) and breast cancers (**Figure** 3B), the clustering of the tumours based on their expression correlation to cell lines did not yield an apparent trend: samples did not group by cancer subtype, stage or grading. Furthermore, even cell lines representing different cancer subtypes displayed a high correlation to all other samples (**Figure** 3A-C). We observed that a small number of tumours displayed consistently lower correlations to all available cell lines. For example, as can be seen in **Figure** 3B for breast samples, one cluster contained six tumours that showed below-median correlations (the median r*_p_* was 0.66) to all the cell lines. This indicates in our view that additional models would need to be established for these samples in order to fully capture the cancer proteome heterogeneity. Equally, it was evident that certain cell lines were proteomic ‘outliers’. For example, three colorectal lines: COCM1, COLO.320DM and RKO, displayed a poorer than average correlation to all tumour samples, with a median r*_p_* of 0.66 (**Figure** 3A).

**Figure 3.**
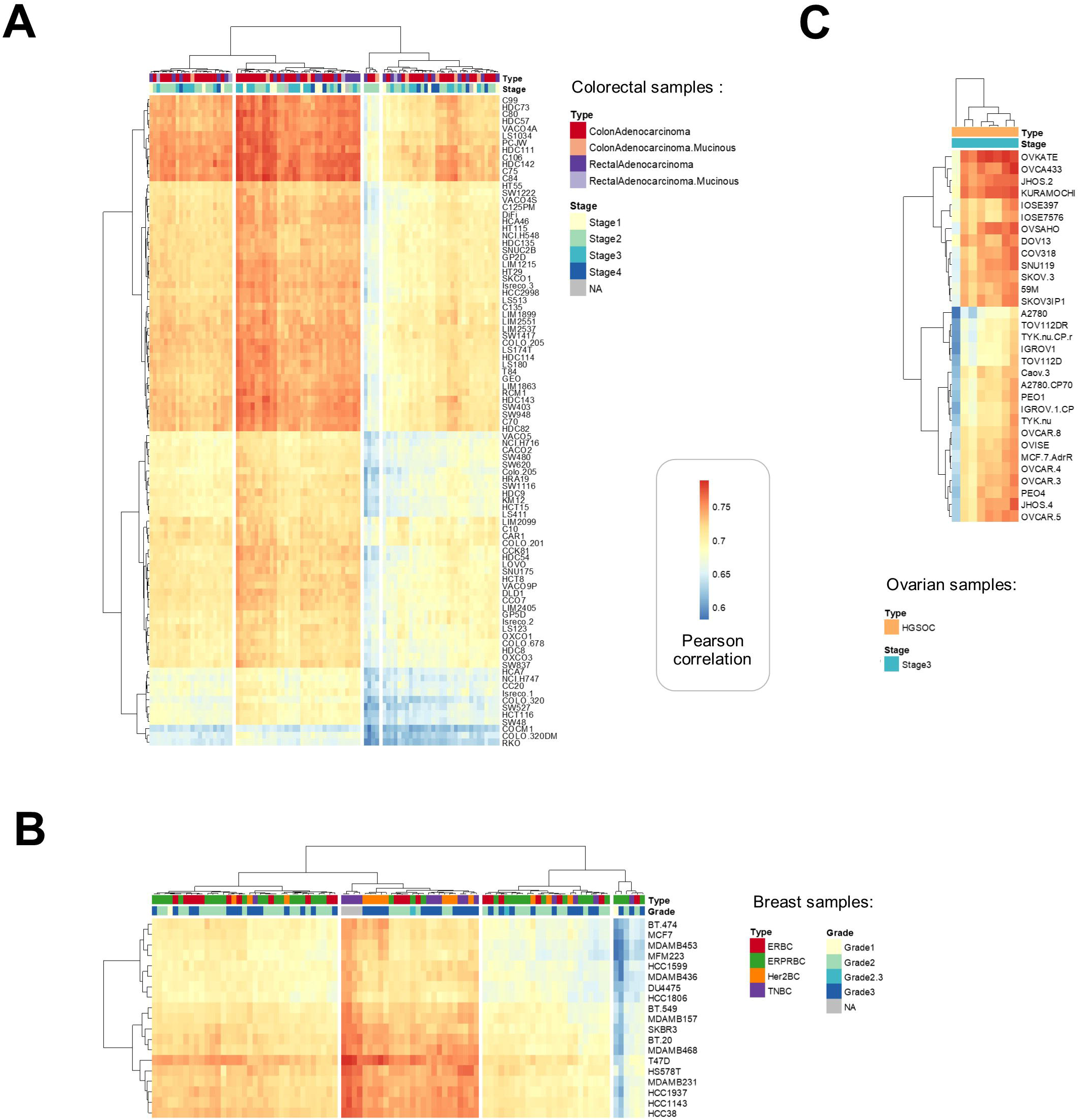
Heatmaps representing correlation matrices between tumour samples (in columns) and cell lines (in rows) for three cancer types: A) colorectal, B) breast and C) ovarian cancer. The protein expression profile of each tumour sample was compared to all cell lines from the corresponding lineage by calculating the Pearson correlation coefficient. This was done across 4,058 proteins overlapping between tumour and cell line samples. Hierarchical clustering was then applied to group the samples. Each cell in the heatmap shows a pairwise comparison and is color-coded according to the calculated correlation value. The small panels on top of a heatmap provide information about the membership of each tumour sample to various categories. A) The top colour bar indicates colorectal tumour type: colon cancer (red) or rectal cancer (purple). The second colour bar is the disease staging used to describe how deeply the primary tumour had grown into the bowel lining. B) The top colour bar shows the breast cancer immunoprofile, and the bottom colour bar indicates the tumour grade, as reported in the original publications. C) In the case of ovarian samples, they were all classified as high-grade serous ovarian cancer and stage 3.

### Cell line-tumour similarity between breast cancer subtypes

Protein expression landscapes appeared to be generally stable and well conserved between the analysed samples. However, despite baseline similarities, it would be possible that specific differences detected between tumour subtypes were not found in cell lines. To assess this, we restricted the following analysis only to breast cancer. This is because this lineage included samples from multiple subtypes in the combined dataset: receptor tyrosine-protein kinase erbB-2 amplified (HER2; 15 tumour and 6 cell line samples), estrogen/ progesterone receptor positive (ERPR; 39 tumour and 10 cell line samples), triple negative (TN, 15 tumour and 31 cell line samples), estrogen receptor positive (ER, 22 tumour samples) and ER/ERPR lymph node metastases (25 tumour samples). Using limma’s linear model^40^, protein expression changes and their statistical significance were calculated between the representative HER2, ERPR and TN cell lines and compared those to changes in tumour tissues (**Figure 4**). In cell lines, 337 differentially expressed gene products were identified in the HER2-TN, 204 in ERPR-TN and 10 in ERPR-HER2 comparison, with a cut-off FDR adjusted p-value of 5%. In tumours, there were 8 differentially expressed gene products in the HER2-TN comparison, 63 in ERPR-TN and 82 proteins in ERPR-HER2. As an example, ERBB2 (UniProt accession number P04626), a tyrosine kinase that is a known marker of HER2 amplified tumours, was overexpressed in both HER2 positive cell lines and tumours (**Figure 4**). However, only five proteins, highlighted as red points in **Figure 4**, had significant protein expression changes in both cell lines and tumours. Whereas statistical significance depends in part on the number of replicates available in each category, a relatively poor agreement in the calculated fold changes between the three subgroups was also apparent. As shown in **Figure 4**, only a small number of proteins displayed similar expression changes in both cell lines and tumours. For example, 38 gene products were overexpressed fourfold (log2 fold change > 2) in ERPR as compared to TN tumours, whereas in the matching cell line comparison, 218 proteins were upregulated fourfold. Interestingly, after performing gene set enrichment analysis (GSEA)^41, 42^ (Methods section) and using the calculated fold changes to pre-rank the proteins, we found that different processes appear to be activated in cell lines and tumours of the same subtype. In case of the ERPR-TN comparison, at a FDR q-value < 0.01 cut-off, only the “hallmark epithelial mesenchymal transition” gene set was enriched among the upregulated proteins in tumours. In cell lines, the enriched sets were “hallmark estrogen response early” and “hallmark estrogen response late”. Five gene sets were enriched among the downregulated proteins in tumours, and15 sets in cell lines, but only two sets were overlapping between the two (“hallmark interferon gamma response”, “hallmark interferon alpha response”). Similar trends were also visible in the two other comparisons (**Supplementary Table 4** contains the summary of all the GSEA results).

**Figure 4.**
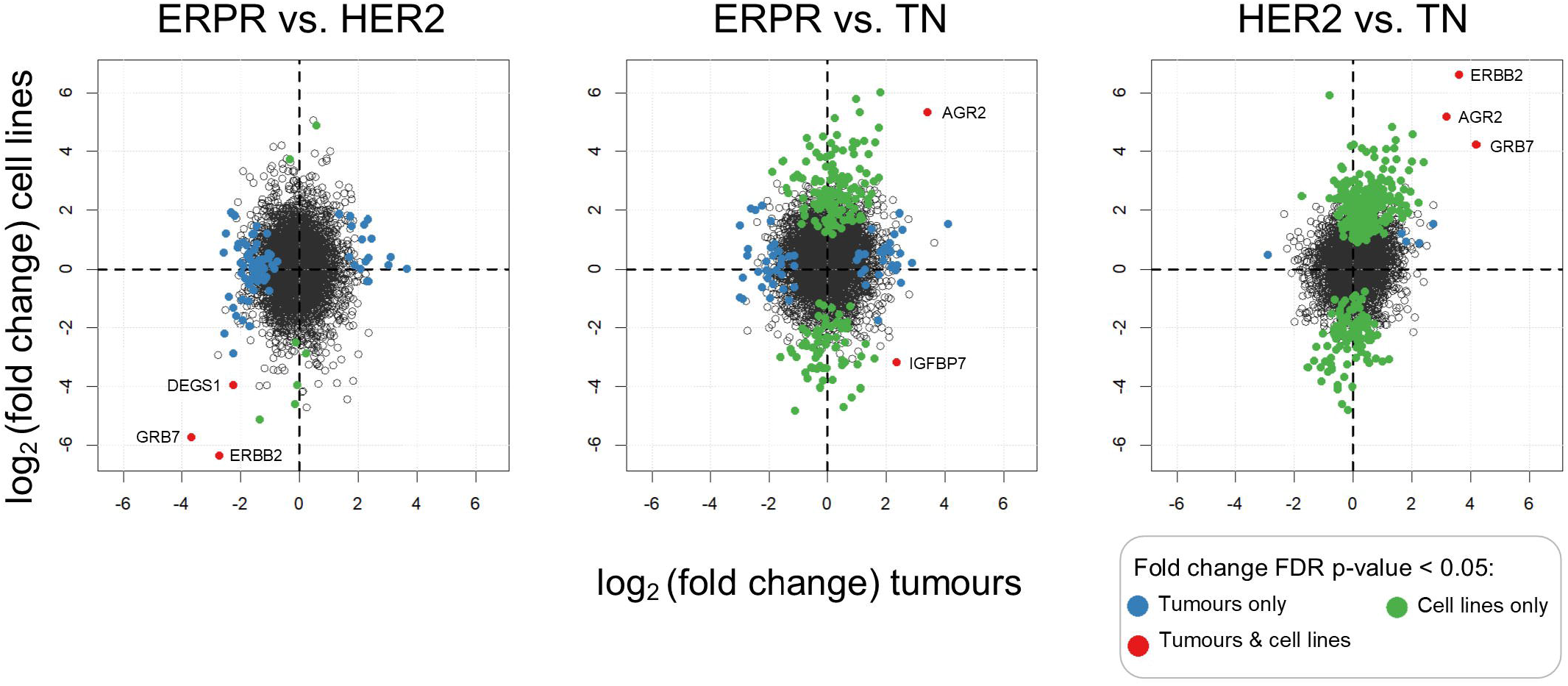
Scatter plots showing the comparison of protein expression changes between breast cancer subtypes (ERPR, HER2 and TN) in cell lines and tumour samples. The x-axis shows the expression changes in tumours and the y-axis the corresponding changes in cell lines. Points are coloured based on the statistical significance of the calculated fold change. Red points represent gene products with statistically significant fold changes in both cell line and tumour comparisons. Blue points represent proteins only statistically significant in tumours and green points only those proteins statistically significant in cell lines.

Overall, this analysis suggests that cell lines provide a good model of baseline protein expression in tumours, as reflected by the high positive correlation found between lineage matching samples. However, specific differences in protein expression, in this case determined only between breast tumour types, appeared to be poorly recapitulated in cell lines.

### Correlation between mRNA and protein expression in cancer cell lines

The correspondence between RNA and protein expression has been previously characterised in cell lines, tumour samples and tissues^43^. In most studies performed in tumour samples so far, the analysis was focused on single cancer types or otherwise, it was limited to a handful of samples for which both mRNA and protein measurements were available. By integrating the aggregated dataset with RNA-seq measurements publicly available already in EA (Methods section), the correlation between mRNA and protein abundance was calculated for 134 cancer cell lines across 13 lineages, including 6,674 gene products that were overlapping between proteomics and transcriptomics data.

### Within sample mRNA-protein expression correlation

To investigate the extent in which mRNA abundances are reflected at the protein level at steady state, the Spearman’s rank correlation coefficient (r*_s_*) was calculated for an average of 6,542 mRNA–protein pairs (some cell lines contained a higher number of mRNA-protein pairs than others), for each of the 134 cell lines. Despite that the original omics measurements were performed in independent studies, all the cell lines displayed a statistically significant (p-value< 2.2 E-16) positive correlation. The median r*_s_* was 0.60 and the values ranged between 0.46 and 0.68 (**Figure** 5A). Box-and-whisker plots were used to show the r*_s_* distributions grouped by lineage (**Figure** 5B). The colorectal (median r*_s_*= 0.61), breast (median r*_s_*= 0.60) and ovarian (median r*_s_* = 0.60) cell lines displayed on average the highest level of correlation, whereas the kidney (median r*_s_* = 0.55), lung (median r*_s_*= 0.56) and blood (median r*_s_*= 0.57) cell lines showed the lowest level of correlation (**Figure** 5B).

**Figure 5.**
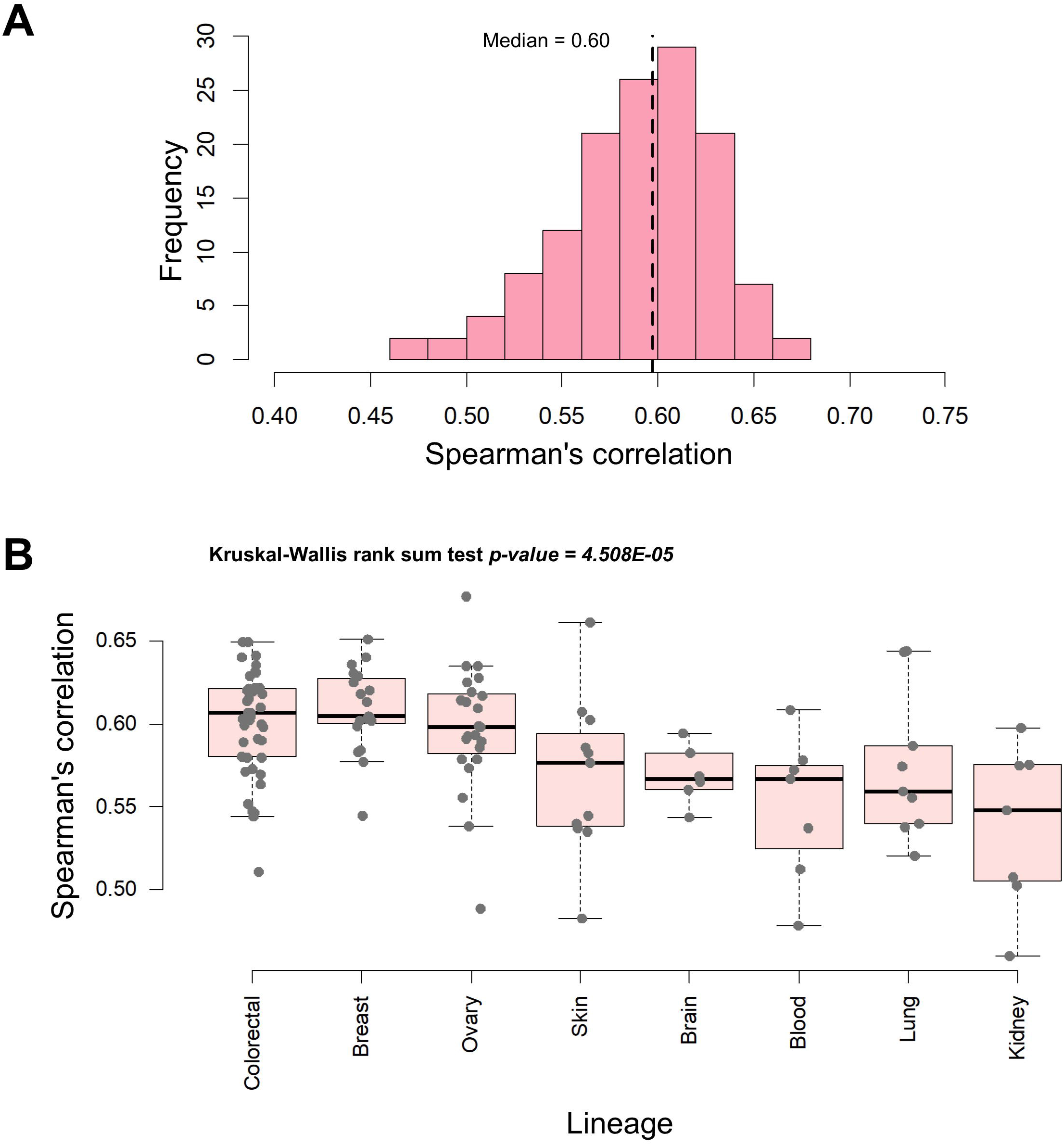
Correlation between mRNA and protein expression levels within the different cell lines. A) Distribution of mRNA-protein correlation using the Spearman correlation in 134 cancer cell lines originating from 13 lineages. The 134 cell lines were those common to this analysis and the existing RNA-seq data, which was obtained from EA. B) Boxplots showing the differences in distribution of mRNA-protein correlations between groups of cell lines originating from various lineages. The box plots show the median (horizontal line), interquartile range (box) and minimum to maximum values of the data. Only lineages with more than three cell lines were included.

**Figure 6.**
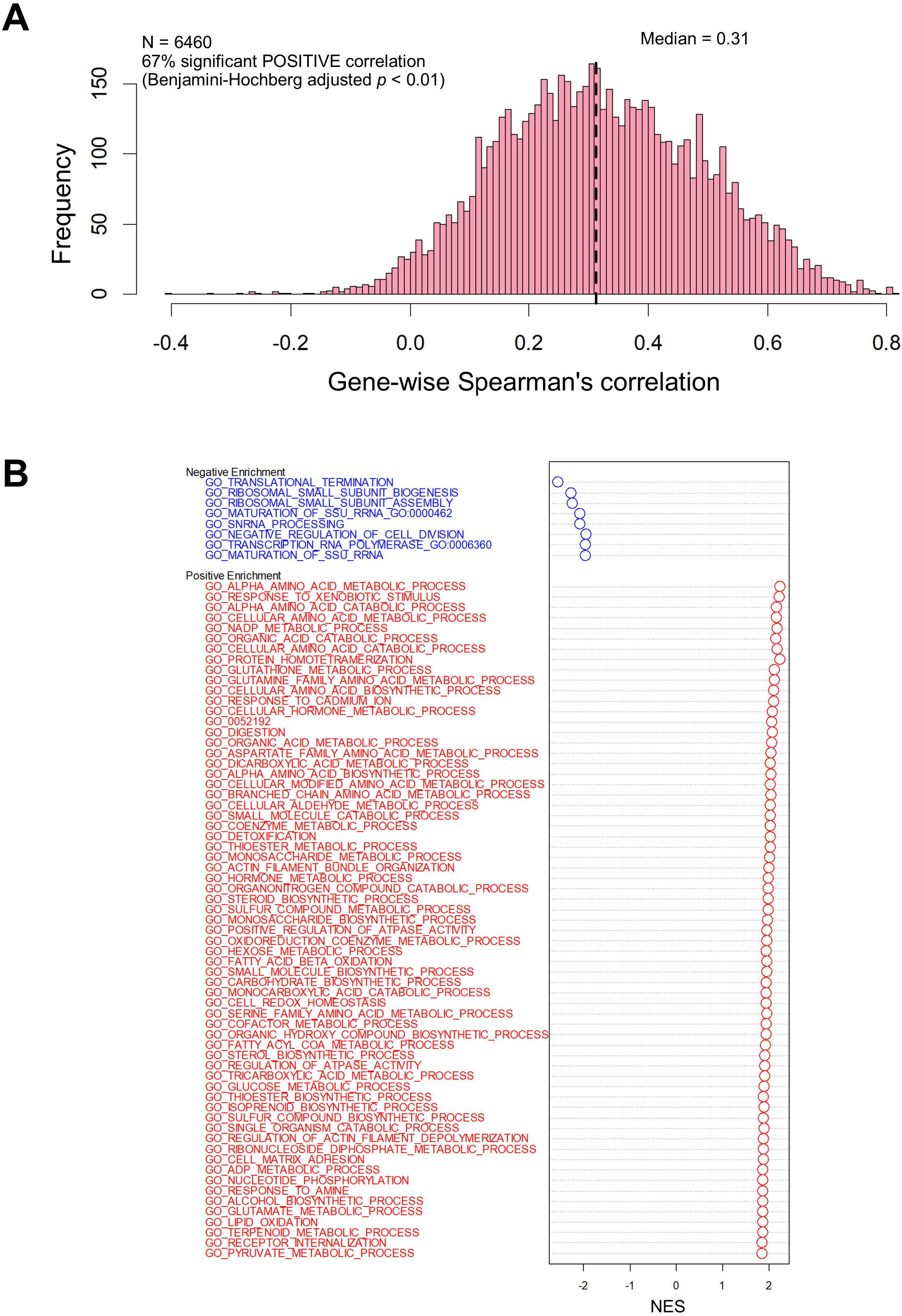
Gene-wise correlation between mRNA and protein levels across all cell lines. A) Histogram of correlation for 6,667 gene-protein pairs across 134 cell lines. B) GSEA showing the top three sets of GO terms significantly represented among those mRNA-protein pairs with a highest (red) and lowest correlation (blue) across the 134 cell lines. Vertical lines indicate the position of the gene set members in the rank ordered list. FDR: False Discovery Rate-corrected p-value.

Albeit small, these differences could have arisen due to random effects or to intrinsic biological factors between the lineage groups. To assess which was the case, a Kruskal–Wallis statistical test was applied, followed by a pairwise Wilcoxon rank-sum post-hoc analysis (**Figure** 5B, Supplementary Table 5). We found that there was a statistically significant difference across lineage groups (Kruskal–Wallis p-value= 4.5 E-05). In addition, four pairwise differences between various cancer types were detected. The most significant one was found between breast-kidney (Benjamini-Hochberg adjusted Wilcoxon p-value= 0.004) and colorectal-kidney (p-value= 0.008) (Supplementary Table 5). Bone, cervix, liver, lymph node and prostate lineages had r*_s_* values close to the overall median (0.60) but were not included in the post-hoc analysis as there were not enough r*_s_* values to calculate a p-value.

Taken together, these results reinforce previous findings where a variable level of agreement between steady state transcript and protein abundance was detected, highlighting the importance of obtaining protein-level measurements in order to gain further insights into a broad range of biological processes.

### Across sample mRNA-protein expression correlation

We investigated the extent of the overall RNA-protein expression correlation across samples. Such analysis consisted on studying how the variation of each transcript and protein originating from the same gene is correlated across all the cell lines. This provided information about whether changes in mRNA levels resulted in abundance changes of the corresponding proteins. An across-sample correlation for a total of 6,667 genes was calculated, of which 4,460 had statistically significant r_s_ values (Benjamini-Hochberg adjusted p-value < 0.01). The median gene-wise r_s_ was 0.31 and the values ranged from −0.40 to 0.82 (**Figure** A). Interestingly, negative correlation values were found only for 2% (160) of the mRNA-protein pairs. However, none of those were significant at a 1% FDR level (Benjamini-Hochberg adjusted p-value). In contrast, 16% (1,065) of genes had a statistically significant correlation that was above 0.5.

Next, the amount of protein variation across cell lines was estimated, by calculating the median abundance and the coefficient of variation for each individual protein. We found that proteins with either high or low variation levels were equally likely to display high correlation between mRNA and protein levels (Supplementary **Figure** 4A). However, the most abundant proteins tended to display higher correlations (Supplementary **Figure** 4B). A possible explanation is that MS experiments are usually biased towards the most abundant proteins, and therefore more accurate measurements of these proteins are obtained.

Additionally, we studied the level of concordance between mRNA and protein variation considering the biological function. GSEA^41, 42^ performed on the list of genes ranked by r_s_, revealed that 65 diverse GO terms were significantly overrepresented among the most correlated mRNA-protein pairs (FDR q-value < 0.01) (Supplementary Table 6). A similar result, where multiple processes were enriched at the top of the ranked list, was also reported for colorectal cancer^18^. The GO terms sets with a largest NES included *protein homotetramerization* (GO:0051289), *alpha-amino acid metabolic process* (GO:1901605), and *response to xenobiotic stimulus* (GO:0009410) (**Figure** B). In contrast, only eight GO term sets, related to the ribosome complex, were overrepresented among the least correlated mRNA-protein pairs. The three sets with lowest NES were: *translational termination* (GO:0006415), *ribosomal small subunit biogenesis* (GO:0042274) and *ribosomal small subunit assembly* (GO:0000028) (**Figure** B, Supplementary Table 6).

## DISCUSSION

Recent large-scale genomics and transcriptomics studies have characterized the molecular diversity of cancer to a great depth. However, in order to understand the relationship between genome and diseased phenotypes, information on protein expression is increasingly relevant. As it is difficult for a single study to cover all proteins of interest or to capture diverse biological conditions (such as different tumour types or disease stages), meta-analysis studies enable the computational integration of multiple studies to provide a combined wide-ranging view.

This study provides a rich resource including an aggregated view of protein expression in cell line and tumour samples, provided to the scientific community through two popular resources: the PRIDE database and EA. Cell lines have indeed provided valuable insights into molecular mechanisms involved in cancer and are generally well-accepted as models of tumour biology. This is possible in part due to the high concordance between molecular signatures, such as RNA expression or single-nucleotide polymorphism (SNP) patterns, which are present both in cell lines and in the tumours of the corresponding lineage^44^. Available studies show different levels of agreement over this statement, and in fact only partial concordance has been found for some of the features^45–47^. Nevertheless, few previous studies have been performed to explore the level of similarity between cell lines and tumours at a proteome-wide level. We have found that the entire proteomes of breast, colorectal and ovarian cell lines generally mirror these coming from the tumours. This was indicated by the relatively high correlations of baseline protein expression profiles. In fact, only a few cell lines displayed low expression correlations. However, the comparison between breast cancer subtypes revealed that specific differences between tumour types are not well represented in the corresponding cell line models and only a small number of proteins showed comparable up or down-regulation in both cases. Furthermore, GSEA showed some biological processes altered between breast cancer subtypes that are different in cell lines and tumours.

It should be pointed out that there are some inherent technical limitations when performing meta-analysis studies like this one. First, although we have used similar quantitative proteomics datasets (only MS1-based quantification approaches performed in Thermo Fischer Scientific Orbitrap instruments), the original data was acquired in different labs in different experimental environments. This inevitably results in the presence of batch effects. We have attempted to remove these and to validate the overall methodology, as described in the Supplementary Methods. Additionally, it has been shown that different batches from the same cell line type can have a higher degree of heterogeneity than what has been generally assumed, as demonstrated recently for HeLa cells^48^. Furthermore, it is known that tumours and cancer cell lines harbour multiple genomic alterations, such as gene fusions or splice variants, which could produce alternative protein sequences. Mutations (e.g. SNPs) can also be acquired as the result of consecutive cell culturing^2, 48^. The MS/MS search strategy used in this study focused only on detecting known coding protein sequences, using the UniProt reference proteome, in the same way as performed in all the original studies. Indeed, cell line-specific genome or transcriptome sequences were not available. Therefore, it was not possible to detect any DNA/RNA sequence changes that could manifest at the protein level. However, the effect of this limitation in the analysis should be small. For instance, in a recent comprehensive study comprising different human tissues^49^, the number of variant peptide sequences detected using matching exome data in the analysis was only a 2.4% (238 out of 9,848 possible amino acid variants).

When examining peptides detected in tumours that were not present in any of the cell lines, we detected signatures enriched in receptor activity regulators as well as in keratinization. Cell lines are purer than tumour samples, which tend to be contaminated with stromal cells. Although, we made every effort to remove common contaminants from the analysis (keratin and others), the ‘tumour-specific’ proteins detected might not necessary reflect endogenous tumour biology. These proteins might have been detected due to tumour immune infiltration^50^, contamination from sample processing and/or contamination from surrounding tissues. Altogether, this highlights some limitations of using *in vitro* cultured cells. While many aspects of protein expression in cancer can be studied using cell lines alone, others (for example, as suggested by our analysis, related to regulation of signalling receptor activity) will likely require better models that are able to model tissue architecture and cell-cell interactions, such as organoid tecnology^51^.

We expect that analogous meta-analysis studies of proteomics datasets will become increasingly popular, due to the unprecedented growth rate of proteomics datasets in the public domain^24^. The availability of these results in widely-used resources such as PRIDE and especially Expression Atlas represent, in our view, the right route for proteomics data to be more accessed and consumed by scientists who are non-expert in proteomics.

## METHODS

### Data sources and curation

Proteomics data from 11 studies (Table 1) was collected from public repositories: PRIDE (https://www.ebi.ac.uk/pride/archive/), MassIVE (https://massive.ucsd.edu/), and CPTAC data portal (https://cptac-data-portal.georgetown.edu/cptacPublic/). Transcriptomics data was obtained from EA (https://www.ebi.ac.uk/gxa/home). The following three RNA-seq experiments, designated as ‘baseline’ in EA, were used in this study: E-MTAB-2706, E-MTAB-2770 and E-MTAB-3983.

Raw proteomics data was manually curated to extract processing parameters, experimental design and sample characteristics. The biological metadata was captured in a Sample and Data Relationship Format (SDRF)^52^ before it was loaded into EA as 11 separate tracks, one for each study. Transcriptomics data have been previously curated in EA, and the biological metadata captured in SDRF format, in a consistent manner with the proteomics data.

### Proteomics data processing

Raw LC-MS data was processed using the MaxQuant software. The MS/MS data was searched in two batches (cell line and tumour data separately) against the UniProt human reference proteome (containing canonical and isoform sequences, download date 31.08.2017, 71,591 sequences) appended with sequences of common contaminants provided by MaxQuant. Search parameters were chosen to reflect those used in the original publications. In all cases, carbamidomethylation of cysteine was set as fixed modification and oxidation of methionine and N-terminal acetylation were set as variable modifications. For studies that used SILAC labelling, appropriate SILAC settings were selected. Enzyme specificity was set to trypsin, LysC, chymotrypsin or GluC (according to the enzymes used in the original study), allowing a maximum of two missed cleavages. MS1 tolerance was set to 10 ppm and MS2 tolerance to 20 ppm for FTMS data and 0.4 Da for ITMS data. PSM (Peptide Spectrum Match), peptide and protein identification FDR was set at 1% at each level. All of the processing parameters are available in the *mqpar-celllines.xml* and *mqpar-tumours.xml* files included in the PRIDE PXD013455 dataset.

### Transcriptomics data processing

Transcriptomics data stored in EA was previously processed using a standardized pipeline^36^. Briefly, the sequencing reads were quality filtered, which involved the removal of adaptor sequences (adaptor trimming), low-quality reads, uncalled bases (e.g. N) and reads arising from bacterial contamination. TopHat2^53^ was used to perform genomics alignment using the reference Ensembl genome (Ensembl release 79). Default TopHat2 parameters were used. The number of reads that mapped to a particular gene (raw counts) were obtained with HTSeq^54^ and normalised gene abundance was calculated as FPKM (fragments per kilobase of exon model per million reads mapped) values^55^. EA data matrices can be downloaded in a tab-delimited format from the corresponding dataset entry. Importantly, for each dataset, technical and biological replicates were averaged and quantile normalised within each set of biological replicates using the limma package^40^. Finally, in cases where a cell line had replicate measurements across datasets, the average FPKM abundance was used.

### Data analysis and integration

All data analyses were performed using custom R scripts. The scripts to generate final quantification file (and selected intermediate files) are available at: https://github.com/J-Andy/Protein-expression-in-human-cancer. MaxQuant outputs various protein quantification metrics such as the summed MS1 intensity, LFQ values, or Intensity Based Absolute Quantification (iBAQ). Here, we have used iBAQ intensities^56^ as a starting point. These were normalised to “parts per billion” (ppb) values. In order to remove batch effects, a procedure to integrate the quantification results was developed and benchmarked as part of this study. The normalisation procedure and various validation results are described in detail in the Supplementary Material in the “Data Processing” section.

### Functional enrichment analysis

GSEA^41, 42^ was carried out using the javaGSEA tool available at http://software.broadinstitute.org/gsea/downloads.jsp. Gene over-representation analysis was performed with online tools GOrilla^37^ and REVIGO^38^. Reactome pathway analysis was performed using the online analysis tool (https://reactome.org/PathwayBrowser/#TOOL=AT) against Reactome version 67. In all cases, p-values were corrected for multiple testing using the Benjamini–Hochberg procedure and the results were considered significant for p-values< 0.01 (unless otherwise stated).

### Data availability

All the results of the study have been made available via the PRIDE database (dataset PXD013455) and in Expression Atlas (E-PROT-19, E-PROT-28, E-PROT-24, E-PROT-20, E-PROT-25, E-PROT-21, E-PROT-22, E-PROT-26, E-PROT-27, E-PROT-18, and E-PROT-23) (Table 1).

The reanalysed public proteomics datasets are indicated in Table 1. The re-used gene expression information coming from cell lines is available in Expression Atlas (accession numbers E-MTAB-2706, E-MTAB-2770 and E-MTAB-3983).

## Supporting information

Supplementary-Table-1-samples-linegae-metadata.txt

## Abbreviations

CCLE: Cancer Cell Line Encyclopedia
CPTAC: Clinical Proteomic Tumour Analysis Consortium
CV: Coefficient of Variation
EA: Expression Atlas
ER: Estrogen Receptor positive (breast cancer subtype)
ERPR: Estrogen/ Progesterone Receptor Positive (breast cancer subtype)
FDR: False Discovery Rate
FPKM: Fragments Per Kilobase of exon model per Million reads mapped
GDSC: Genomics of Drug Sensitivity in Cancer
GO: Gene Ontology
GSEA: Gene Set Enrichment Analysis
HER2: Receptor tyrosine-protein kinase erbB-2 amplified (breast cancer subtype)
iBAQ: Intensity Based Absolute Quantification
LFQ: Label Free Quantification
MS: Mass Spectrometry
NES: Normalised Enrichment Score
NGS: Next-generation sequencing
ppb: parts per billion normalisation
PSM: Peptide Spectrum Match
RPPA: Reverse-Phase Protein Arrays
SDRF: Sample and Data Relationship Format
TCGA: The Cancer Genome Atlas
TN: Triple Negative (breast cancer subtype)Abbreviations

## Acknowledgements

The authors would like to thank Dr Pedro Beltrao for valuable discussions and critical feedback.

The authors also want to express their gratitude to all the original authors of the deposited public datasets used in this study.

This study has been funded by the Wellcome Trust [grant number 208391/Z/17/Z] and by EMBL core funding.

## SUPPLEMENTARY MATERIAL

### SUPPLEMENTARY FIGURES

**Supplementary Figure 1.**
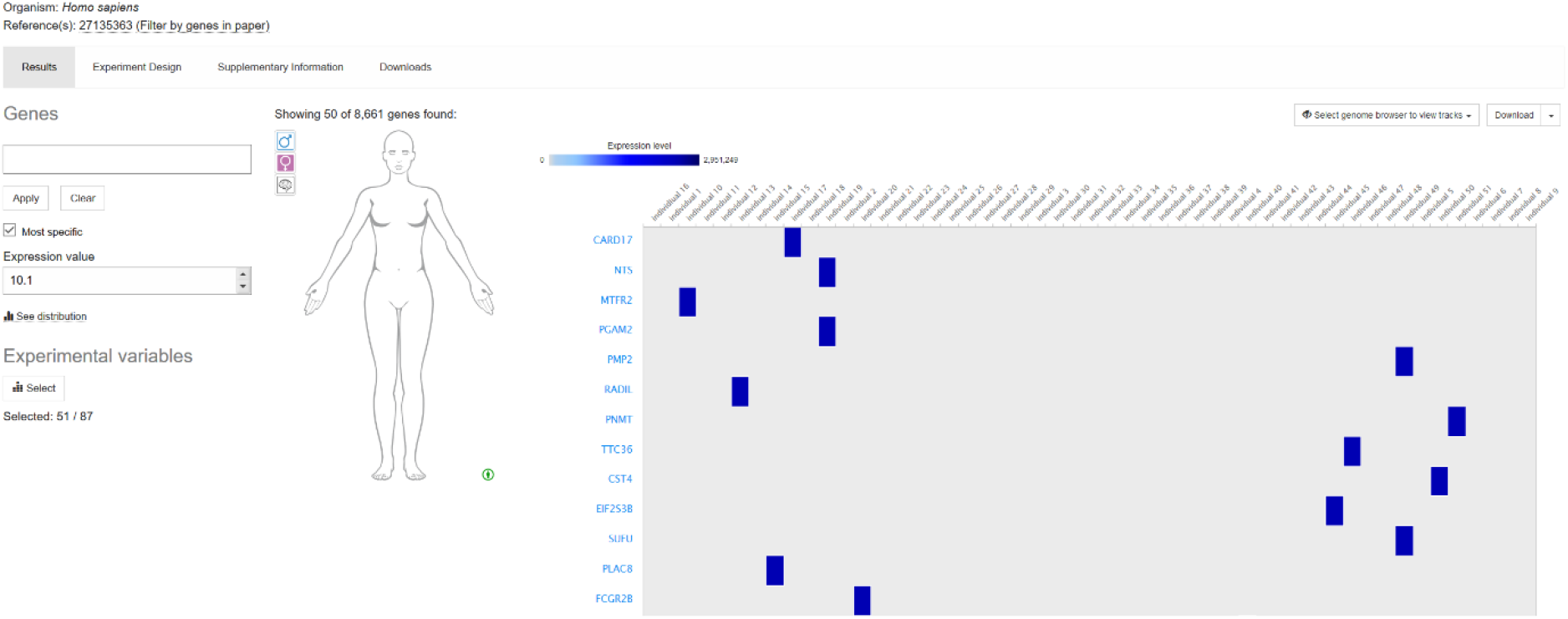
Baseline protein expression view in Expression Atlas. Screenshot containing an example of the baseline expression view for dataset E-PROT-26 (“Deep proteomic profiling of luminal breast cancer progression”, https://www.ebi.ac.uk/gxa/experiments/E-PROT-26/Results) within Expression Atlas. Protein expression levels are displayed in a heatmap.

**Supplementary Figure 2.**
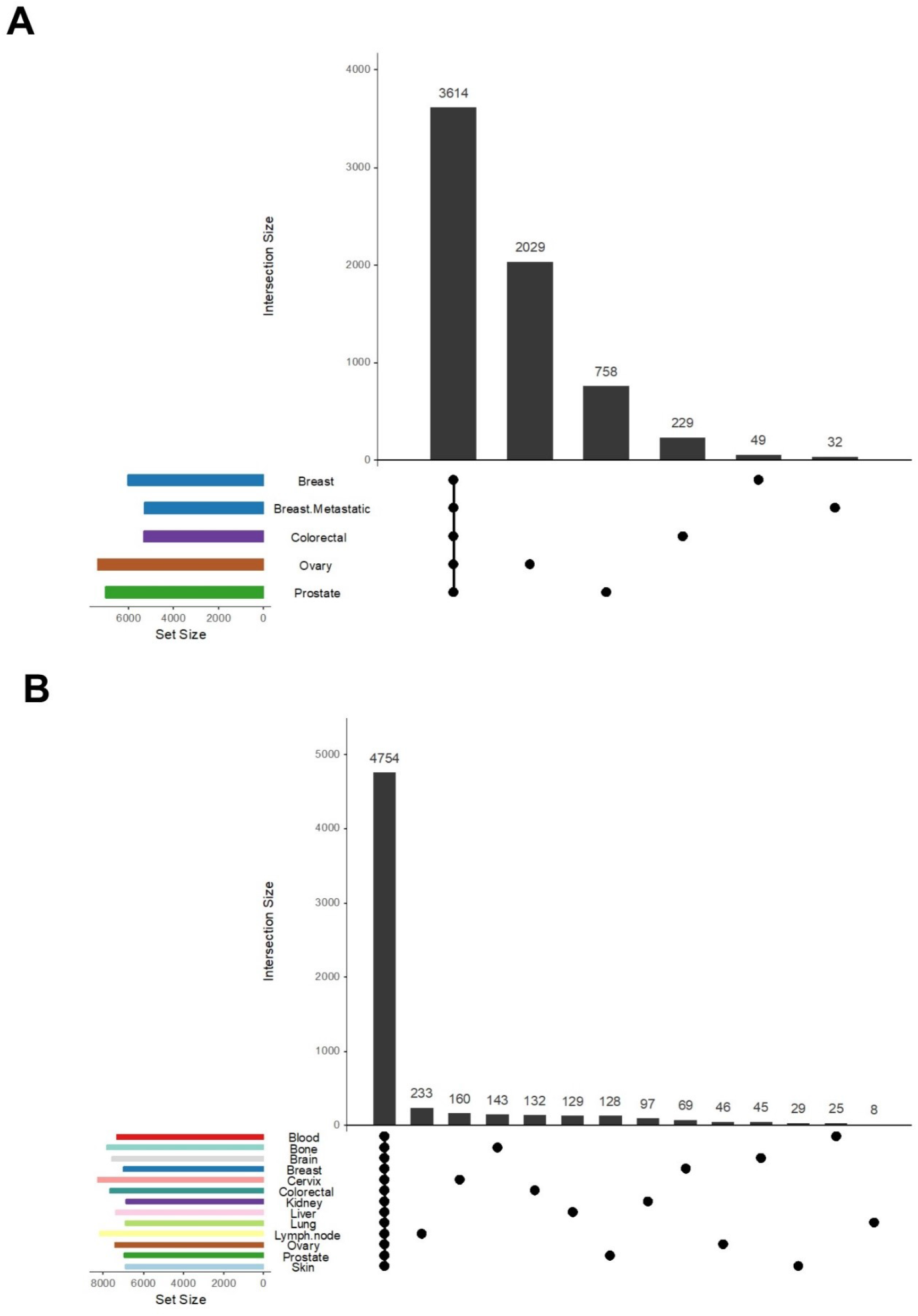
Upset plot summarising proteins detected across cancer types. Counts of proteins detected in the majority (meaning ≥ 50% of samples) of A) tumour samples; and B) cell lines. Black dots indicate the individual sample or combinations of samples for which the number of proteins unique to that combination is displayed. This plot was created with the ‘UpSetR’ R package^1^.

**Supplementary Figure 3.**
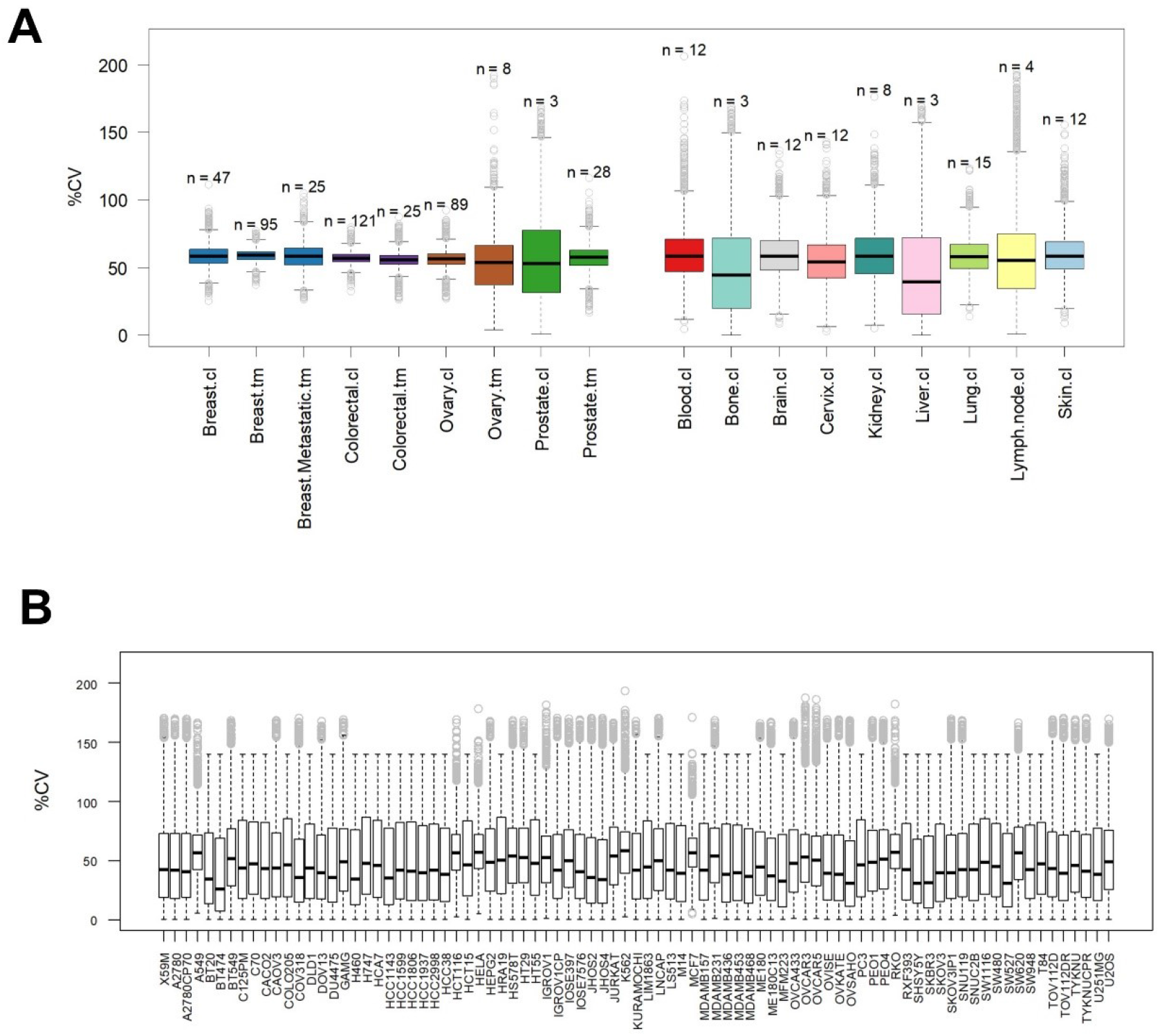
Variability of protein expression in cell lines and tumours. Variation in protein expression across cancer samples. A) Boxplots show the distribution of the Coefficient of Variation (CV) metric for all proteins, stratified by cancer lineage (different colours) and separated into cell line (.cl) or tumour (.tm) samples. The number of samples per type (n) is also indicated. B) Boxplots show the distribution of the CV metric across biological replicates in cell lines. Only cell lines with at least 3 replicates were included.

**Supplementary Figure 4.**
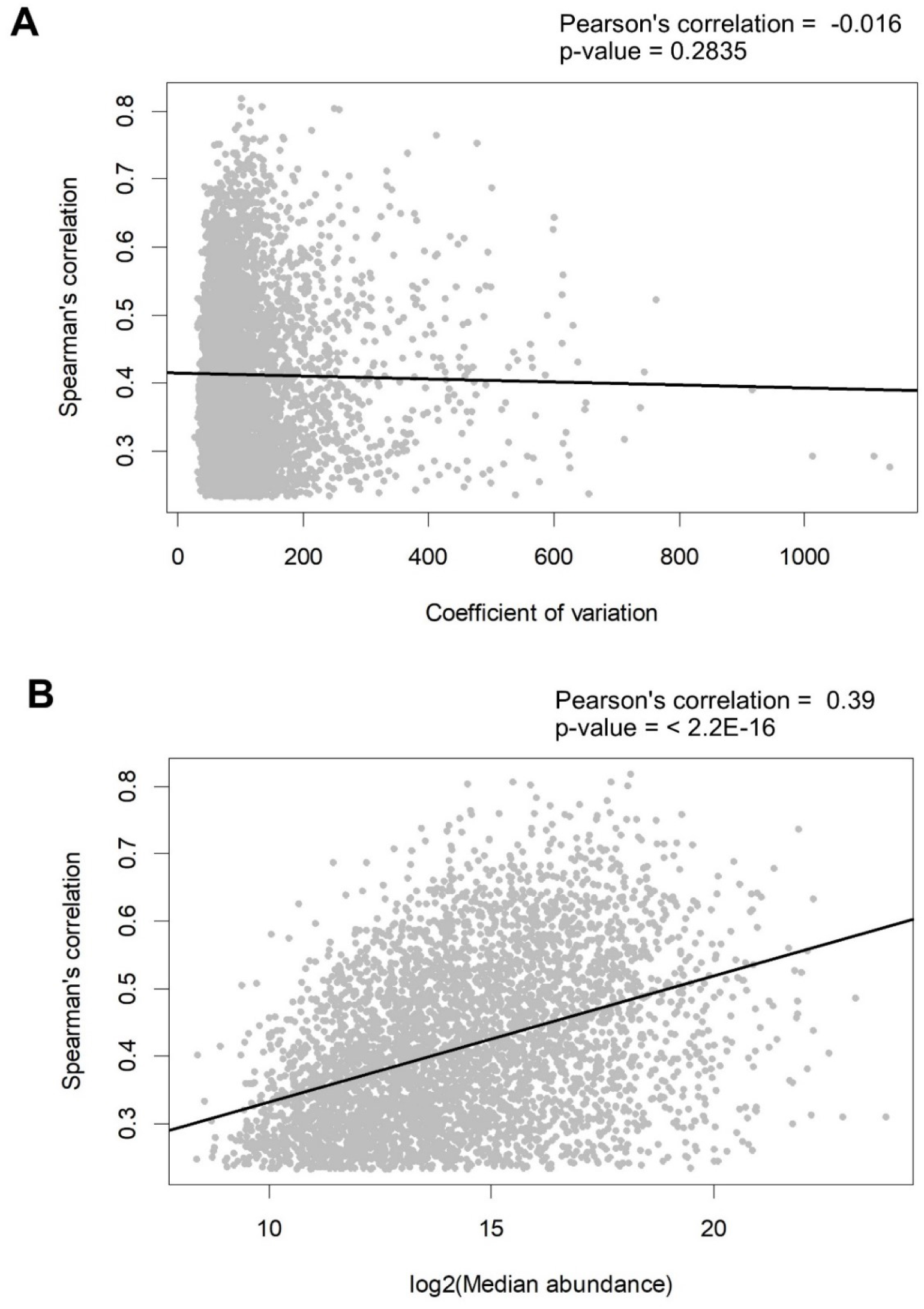
Dependence of mRNA-protein correlation on protein variation and abundance. Scatter plots show the extent in which mRNA-protein correlation is associated with A) the variation in protein abundance across cell lines; and B) the average protein abundance, expressed as parts per billion-normalised iBAQ (Intensity Based Absolute Quantification) values across cell lines. In panel A, it can be observed that an increase in protein abundance variation is not correlated with mRNA-protein covariation across the cell lines. In panel B, mRNA-protein covariation tends to increase in parallel with protein abundance.

**Supplementary Figure 5.**
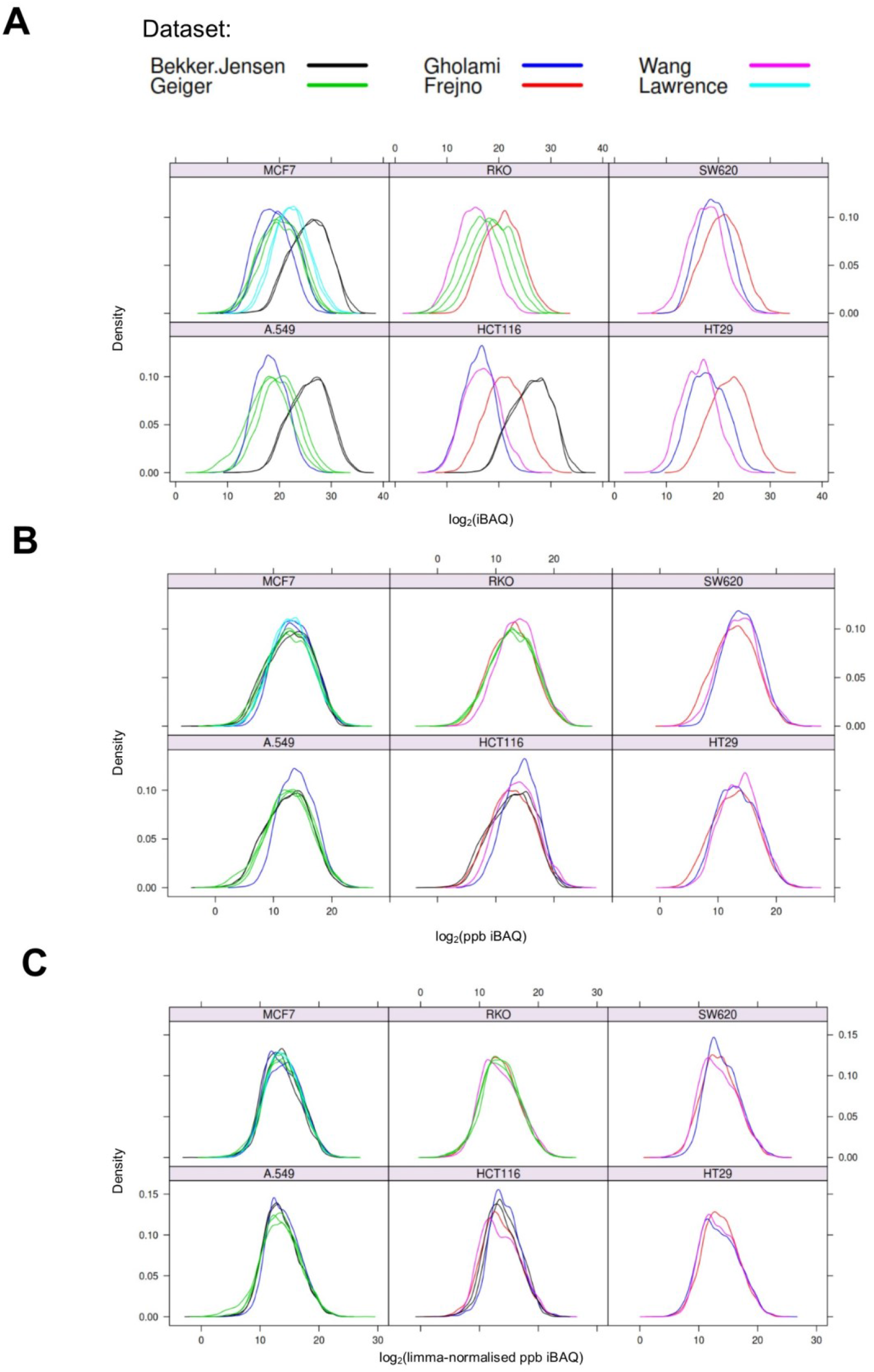
Density plots showing the distributions of log2 transformed iBAQ intensities in six cell lines acquired in at least three studies. Density distribution plots of protein expression values in six cell lines for which biological replicates (in a minimum of three independent studies) were available. Panel A) shows the un-normalised iBAQ values. Panel B) shows the ‘ppb’ normalised iBAQ values. Panel C) shows the final quantification values after applying missing data imputation and batch effects removal with limma. Different colours indicate the study of origin. The plots reveal that strong global effects are present due to the study of origin.

**Supplementary Figure 6.**
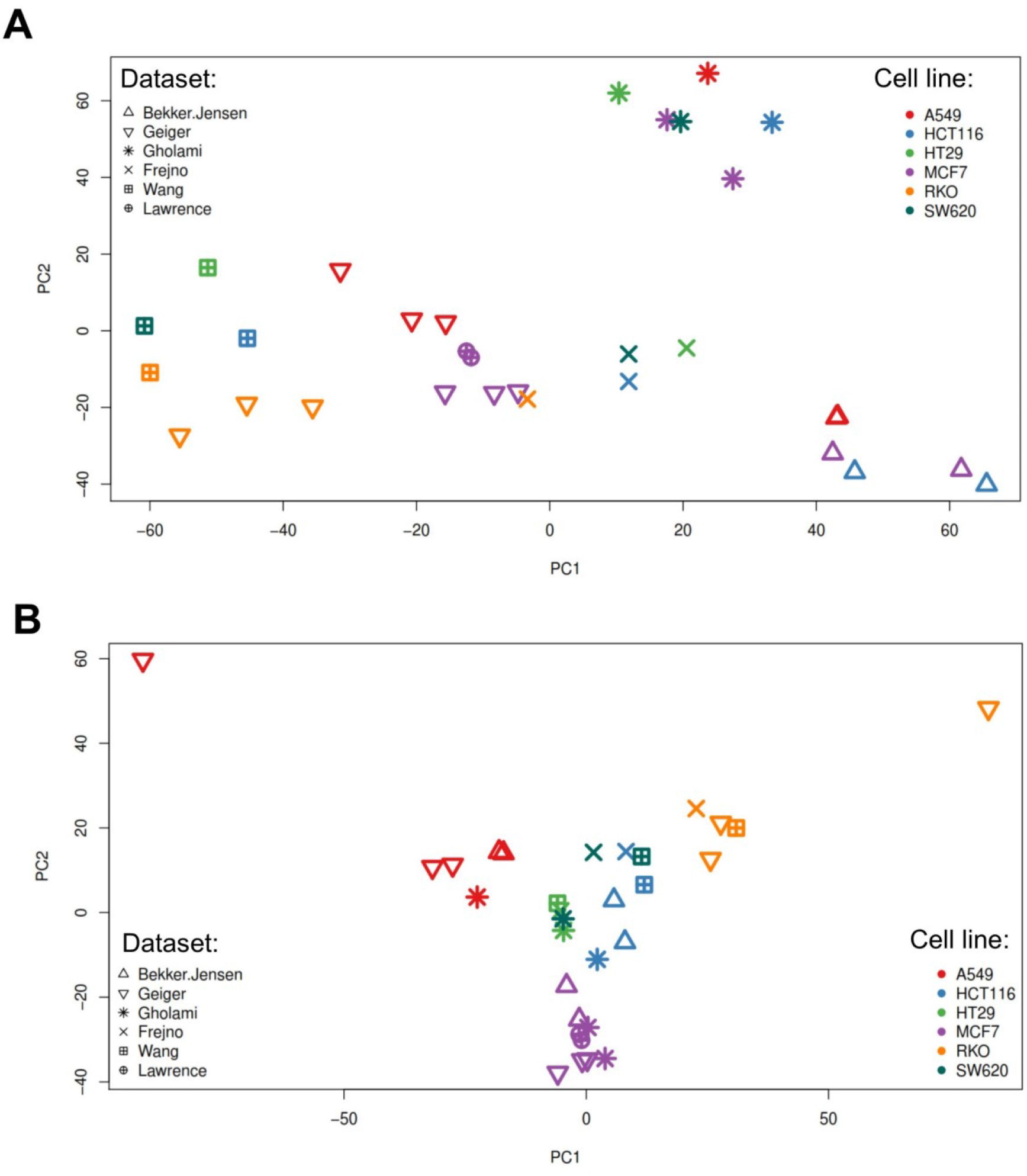
Principal component analysis. A complete matrix across six cell lines (A549, HCT116, HT29, MCF7, RKO and SW620) was used for PCA analysis. This included 2,914 protein expression values. The first two principal components explaining 35% and 30% of the variance are displayed. The colour of the points indicates the sample types whereas the shape of the point indicates the corresponding study. Panel A) shows the PCA space after “ppb” normalisation. It can be observed that points (individual samples) cluster according to the study of origin. Panel B) shows the PCA space after limma batch effects removal. It can be observed that cell lines cluster together based on the lineage rather than based on the study of origin.

### SUPPLEMENTARY METHODS

#### 1. Data Processing

Overall, 7,171 MS (Mass Spectrometry) proteomics runs coming from 11 large-scale quantitative cancer related proteomics studies were selected, manually curated and re-annotated after performing a comprehensive review of the current literature and the public availability of the corresponding datasets. These studies were selected since they all employed a similar MS platform (Thermo Fisher Scientific Orbitrap) and the resulting protein quantification could be based on the intensity of the peptide precursor ions (MS-1 based quantification^2^). The final set of samples was assembled in September 2018 and is summarised in Table 1 in the main text. The original raw data files were reanalysed using MaxQuant^3^ (MQ) as indicated in the main text (see also **Figure** 1A there). For data processing, the minimum peptide length was set to seven amino acids. Unique and razor peptides, as well as peptides containing Oxidation (in M) and Acetyl (in the protein N-term) modifications were used for quantification purposes. All of the used processing parameters are available in the *mqpar-celllines.xml* and *mqpar-tumours.xml* files included in the PRIDE dataset identifier PXD013455.

Since quantitative proteomics data originating from different studies is heterogeneous and likely to contain batch effects, we developed and benchmarked a procedure to integrate the quantification results. The procedure is described below.

##### i. Selection of protein quantification values

iBAQ protein quantification values were obtained from the corresponding MQ *proteinGroups.txt* files. In some studies, multiple digestion enzymes were employed to characterize the same sample, for example in the dataset from ref^4^ (see Table 1 in the main text), where all cell line samples were digested with both trypsin and LysC. Because iBAQ quantification takes into account all theoretically observable peptides in a given protein, and these will differ depending on the proteolytic enzyme used, the quantitative analysis was limited to tryptic-digested samples only.

##### ii. Data normalisation

Cell line-derived quantitative data was used to develop and benchmark a normalisation procedure. This was possible because six cell lines (A549, HCT116, HT29, MCF7, RKO and SW620) in the aggregated dataset were acquired in at least three independent studies. It was then assumed that the highest amount of variability in protein expression in those cell lines, and indeed in any other sample type, should arise due to biological differences (i.e. different sample origin), rather than due to technical artefacts, such as the study of origin.

First, the presence of batch effects in the reprocessed data was evaluated by plotting density distributions of log2 transformed iBAQ intensities (Supplementary Figure 5A). Upon visual examination of the plots, it was evident that global biases were present between different studies. To correct for these, a two-step normalisation process was applied. As a first step, the individual iBAQ intensities were transformed to “parts per billion” (ppb) for each of the MS runs. Each protein iBAQ intensity value was scaled to the total amount of signal in a given MS run and transformed to ppb, as expressed in the following equation:

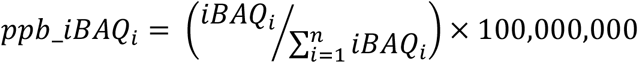

As seen in Supplementary Figure 5B this procedure mostly removed global differences in the distribution of protein abundances between the studies, which can occur due to different amounts of protein loaded on the chromatographic column, or simply because different MS instruments record data on different numerical scales, among other reasons. However, examination of a principal component analysis (PCA) plot, based on 2,914 proteins quantified in all of the six cell lines, suggested that this simple scaling procedure did not remove the main batch effects due to study of origin (Supplementary Figure 6A).

The cell line and tumour datasets were then filtered to include only proteins that were present in at least 50% of all assays (MS runs). This resulted in the cell line dataset containing 6,514 proteins and a 14% of missing values, and the tumour dataset containing 5,363 proteins and a 16% of missing values. Missing values were then imputed using the conventional Singular Value Decomposition method implemented in the pcaMethods R package (“SVDimpute” function^5^). Finally, the main batch effects were removed separately for each dataset (“cell lines” and “tumours”) using the limma R package^6^ including the study of origin as a covariate (“removeBatchEffect” function). Post-normalisation PCA analysis (Supplementary Figure 6B) and inspection of density distribution plots (Supplementary Figure 5C) confirmed that the large batch effects among studies of different origin were removed. In the last step, the two datasets were merged by cross-referencing the leading razor protein identifiers.

It must be emphasised that it was only possible to correct for batch effects within each individual “cell lines” or “tumours” protein expression matrix. That is because in many cases, measurements from the same biological sample (i.e. distinct cell line or same tumour type) were acquired in multiple batches (i.e. studies). However, very few overlapping samples were acquired among the “cell lines” and “tumours” datasets.

## Abbreviations

iBAQ: Intensity Based Absolute Quantification
CV: Coefficient of Variation
MS: Mass Spectrometry
MQ: MaxQuant
PCA: Principal Component Analysis

